# Comprehensive analysis of lncRNAs reveals candidate prognostic biomarkers in multiple cancer types

**DOI:** 10.1101/861039

**Authors:** Keren Isaev, Lingyan Jiang, Christian A. Lee, Ricky Tsai, Fiona Coutinho, Peter B. Dirks, Daniel Schramek, Jüri Reimand

## Abstract

Long non-coding RNAs (lncRNAs) are increasingly recognized as functional units in cancer pathways and powerful molecular biomarkers, however most lncRNAs remain uncharacterized. Here we performed a systematic discovery of prognostic lncRNAs in 9,326 patient tumors of 29 types using a proportional-hazards elastic net machine-learning framework. lncRNAs showed highly tissue-specific transcript abundance patterns. We identified 179 prognostic lncRNAs whose abundance correlated with patient risk and improved the performance of common clinical variables and molecular tumor subtypes. Pathway analysis revealed a large diversity of the high-risk tumors stratified by lncRNAs and suggested their functional associations. In lower-grade gliomas, discrete activation of HOXA10-AS indicated poor patient prognosis, neurodevelopmental pathway activation and a transcriptomic similarity to glioblastomas. HOXA10-AS knockdown in patient-derived glioblastoma cells caused decreased cell proliferation and deregulation of glioma driver genes and proliferation pathways. Our study underlines the pan-cancer potential of the non-coding transcriptome for developing molecular biomarkers and innovative therapeutic strategies.

## INTRODUCTION

The human genome encodes numerous long non-coding RNAs (lncRNAs) that lack protein-coding potential and are sparsely annotated [1, 2]. A recent survey annotated nearly 20,000 high-confidence human lncRNA genes of at least 200 nucleotides in length, indicating that lncRNAs are at least as common as protein-coding genes [2]. Globally, lncRNAs are transcribed at lower levels compared to protein-coding genes and exhibit transcript abundance patterns specific to tissue types and developmental stages [1, 3]. lncRNAs are involved in the regulation of cellular processes through multifunctional interactions with the genome, transcriptome and proteome [4, 5]. Individual lncRNAs are increasingly recognized as key players in diverse biological processes such as chromatin remodeling in X chromosome inactivation [6], post-transcriptional gene regulation through alternative splicing [7], and epigenetic silencing through histone modification [8]. Computational analysis of lncRNAs enables systematic functional insights and gene prioritization. For example, k-mer analysis identified non-linear sequence similarities between lncRNAs that were informative of protein-RNA interactions and sub-cellular localization [9]. However, the vast majority of lncRNAs lack functional annotations and most of our knowledge of non-coding genes is based on a few well-studied examples.

lncRNAs are increasingly implicated in cancer hallmark pathways such as proliferation, angiogenesis, growth suppression, cell motility and immortality [10]. Specific well-studied lncRNAs are now recognized as biomarkers for diagnosis, prognosis and therapy of cancer. The first lncRNA-based biomarker gene *PCA3* is specifically expressed in prostate cancer tissue relative to normal prostate tissue [11] and is now used in non-invasive tests that complement standard serum-based tests of prostate-specific antigen [12]. The lncRNA *HOTAIR* is involved in cancer progression and metastasis through chromatin remodeling and its increased transcript abundance in breast cancer is a robust predictor of tumor metastasis and patient survival [13]. Transcriptional profiling of normal and tumor samples has revealed numerous tissue-specific lncRNAs [1, 15, 16], indicating further potential for discovery and development of cancer biomarkers based on the noncoding transcriptome. Some lncRNAs are also frequently mutated in cancer genomes and recent studies have identified candidate driver mutations by surveying whole-genome sequencing data in multiple cancer types [17, 18]. Projects such as The Cancer Genome Atlas (TCGA) [19], International Cancer Genome Consortium (ICGC) [20], METABRIC [21] and others have accumulated multi-omics datasets and patient clinical profiles for thousands of cancer samples. These resources have enabled biomarker studies that associated cancer patient prognosis with transcript abundance of protein-coding genes and their genetic and epigenetic alterations [22-25]. However, associations of lncRNAs with cancer patient survival and biological function remain largely unexplored. A recent study characterized recurrent hypomethylation patterns affecting a thousand lncRNAs in the TCGA PanCanAtlas cohort and identified the *EPIC1* lncRNA as a marker of poor prognosis in a subset of breast cancers [26]. Another TCGA study associated mutations and transcript abundance profiles of lncRNAs with regulatory networks and molecular pathways and nominated candidate oncogenic and tumor suppressive lncRNAs, some of which were functionally validated in cancer cell lines [27]. Analysis of cell-cycle correlated lncRNAs revealed a subset of S-phase enriched lncRNAs whose transcript abundance profiles correlated with patient survival in multiple TCGA cohorts [28]. However, those studies did not analyze robust prognostic performance of lncRNAs using machine-learning and cross-validation approaches, indicating further potential to systematically discover lncRNAs as candidate prognostic biomarkers of multiple cancer types.

Here we evaluated the transcript abundance profiles of nearly 6,000 lncRNAs as prognostic biomarkers in human cancers. Using a comprehensive machine-learning analysis, we compiled a robust catalogue of prognostic lncRNAs across nearly 10,000 tumors of 29 types from the TCGA PanCanAtlas project [22, 29]. The majority of our candidate lncRNAs showed improved prognostic potential compared to standard clinical features and molecular tumor subtypes. We associated prognostic lncRNAs with large-scale deregulation of hallmark cancer pathways, revealing extensive functional diversity of high-risk tumors and potential roles of lncRNAs. Using functional experiments in patient-derived glioma cell lines, we show that knockdown of the lncRNA *HOXA10-AS* led to reduced cellular proliferation and transcriptional de-regulation of hallmark cancer pathways and driver genes. Our study highlights the translational utility of the human non-coding transcriptome for cancer biomarker discovery and provides a catalogue of high-confidence lncRNAs for functional experiments and biomarker studies.

## RESULTS

### Long non-coding RNAs (lncRNAs) show tissue-specific transcript abundance and patient survival associations in multiple cancer types

We first characterized the transcript abundance of lncRNAs across 9,326 patients from 29 cancer types with matched RNA-sequencing (RNA-seq) data and clinical annotations of the TCGA PanCanAtlas dataset [22, 29] (**Supplementary Table 1**). We identified 5,785 high-confidence lncRNAs that were annotated by both the FANTOM CAT project [2] and the Ensembl database [30] (**Supplementary Table 2**). We first asked whether the lncRNAs showed tissue-specific transcript abundance patterns in the TCGA pan-cancer dataset. Unsupervised clustering of lncRNA transcriptomes using the UMAP dimensionality reduction algorithm [31] revealed a robust grouping of tumor samples by organ systems and histological subtypes (**Figure 1A**), akin to multi-omics data of protein-coding genes [29]. For example, the clusters indicated lncRNA-based transcriptional similarity of lower-grade gliomas and glioblastomas of the brain (LGG, GBM), colon and rectum adenocarcinomas (COAD, READ), and four types of squamous carcinomas (BLCA, LUSC, HNSC, CESC). Highly tissue-specific lncRNA abundance patterns suggest that the non-coding transcriptome includes uncharacterized diagnostic and prognostic biomarkers.

**Figure 1.**
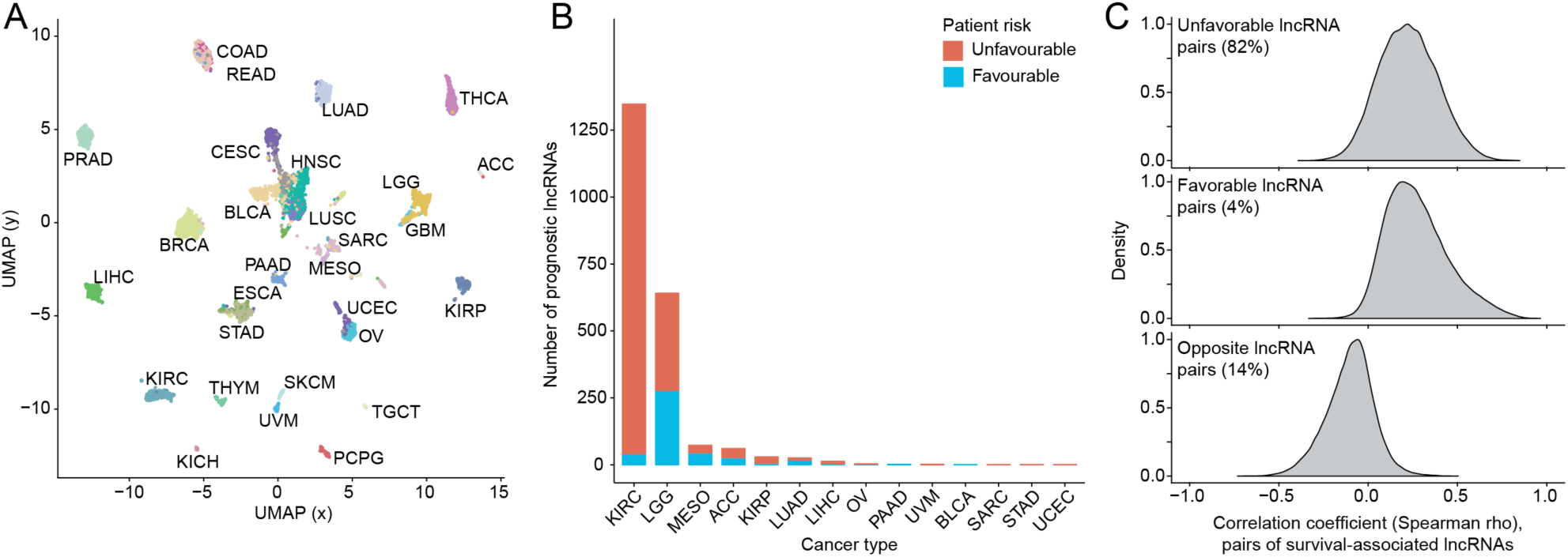
Tissue specificity and patient survival associations of IncRNAs in multiple cancer types. **A.** Unsupervised clustering of IncRNA transcript abundance across 29 cancer types in TCGA indicates high tissue-specificity of IncRNA transcription. **B.** Thousands of individual IncRNAs are significantly associated with overall patient survival in multiple cancer types (Cox PH, *FDR* < 0.05). **C**. Survival-associated IncRNAs are characterized by highly redundant transcript abundance profiles. Density plots show correlation coefficients from an exhaustive pair-wise analysis of all survival-associated IncRNAs. IncRNA pairs with matching risk profiles (both unfavourable, top; both favourable, middle) are often positively correlated while IncRNA pairs with opposing risk profiles are often negatively correlated in transcript abundance. Thus the non-coding transcriptome represents a redundant space for prognostic marker discovery that is confounded by gene regulatory and clinical features of tumors.

As a pilot study of lncRNAs as prognostic markers in human cancers, we associated lncRNA transcript abundance with overall patient survival using Cox proportional-hazards (PH) models. We used individual lncRNAs as predictors in combination with standard clinical variables such as patient age, sex, tumor stage and/or grade available in TCGA. Nearly half of lncRNAs were significantly associated with overall patient survival in at least one cancer type (2,740 of 5,785, 47%, Wald test, *FDR* < 0.05), with the majority of lncRNAs found in kidney renal cell carcinoma (KIRC) and lower-grade glioma (LGG) (**Figure 1B**). Most of these lncRNAs were associated with survival in only one cancer type (2,203/2,740 or 80%), confirming tissue-specificity of lncRNA transcription. The majority of lncRNAs appeared hazardous (81%) as their transcript abundance was associated with poor prognosis. Interestingly, 18% of lncRNAs were zero-dichotomized based on their discrete transcriptional activation patterns, as one group of patients showed high transcript abundance of a given lncRNA while the other patient group showed complete lncRNA silencing. These characteristics suggest a high potential for biomarker discovery in non-coding cancer transcriptomes.

Having identified thousands of survival-associated lncRNAs in the pilot analysis, we asked whether these represented robust and independent signals of transcript abundance. We performed an exhaustive co-expression analysis of all 1,116,955 pairs of survival-correlated lncRNAs in their corresponding cancer types and found that a large fraction (35%) were significantly correlated in transcript abundance (Spearman correlation, rho > ±0.3 and *FDR* < 0.05; **Figure 1C**). As expected, lncRNA pairs with matching prognostic risk were often positively correlated while pairs of lncRNAs with opposing risk correlated negatively. Thus, this large pool of putatively survival-associated lncRNAs represent a considerably narrower space of transcriptional signatures that are confounded by factors such as epigenetic or transcriptional co-regulation, patient clinical characteristics and tumor subtypes. This analysis indicates that many lncRNAs are expected to be transcriptionally correlated with patient survival in statistical tests however their confounders and high rate of co-expression limit their use in prognostic models designed to evaluate previously unseen patients. A systematic computational strategy is needed to distinguish representative and robust lncRNAs as prognostic biomarkers.

### Elastic net proportional-hazards framework identifies 179 prognostic lncRNAs

To identify robust and non-redundant prognostic lncRNAs, we implemented a machine-learning strategy of Cox-PH models with elastic net regularization by adapting earlier studies on the prognostic evaluation of omics data [25, 32] (**Supplementary Figure 1**). Briefly, multivariate regression models with high-confidence lncRNAs as predictors and patient overall survival as response were fitted separately for each cancer type across 1,000 cross-validations with 70/30% data splits for training and testing. Each model initially included a pool of nominally survival-associated lncRNAs for the given cancer type that were evaluated based on training data (Cox PH *P* < 0.05). The subsequent feature selection step extracted a subset of lncRNAs as high-confidence predictors for that cross-validation iteration. These multivariate models were then evaluated on test data using the concordance index (c-index), an accuracy measure for risk models with censored survival data [33]. We also fitted baseline models as controls that included only clinical variables as predictors (*e.g.*, tumor stage, grade, patient age and sex, as available in TCGA), and additional combined models that included as predictors both the set of clinical variables and all pre-selected transcript abundance profiles of lncRNAs. We evaluated the entire series of multivariate lncRNA-based survival models trained through cross-validations.

Prognostic models of lncRNA-based predictors showed consistently superior performance in terms of concordance index values in nine cancer types, compared to baseline models that only included clinical variables (Wilcoxon rank-sum test, *FDR* < 0.05; **Supplementary Figure 2**). Combining clinical variables and lncRNA transcript abundance profiles as predictors further improved prognostic performance of our models in 12 of 28 cancer types. To evaluate false-positive rates of our strategy, we also generated 100 simulated datasets for each cancer type by randomly reassigning patient survival data within each cohort of a specific cancer type. As expected, c-indices from the simulated datasets were consistently lower than those obtained from true data and centered on the performance value of a random predictor (c = 0.5), lending confidence to our strategy (**Supplementary Figure 3**). These observations underline the added value of analyzing lncRNAs as prognostic biomarkers and suggest follow-up validation analyses in additional patient cohorts.

**Figure 2.**
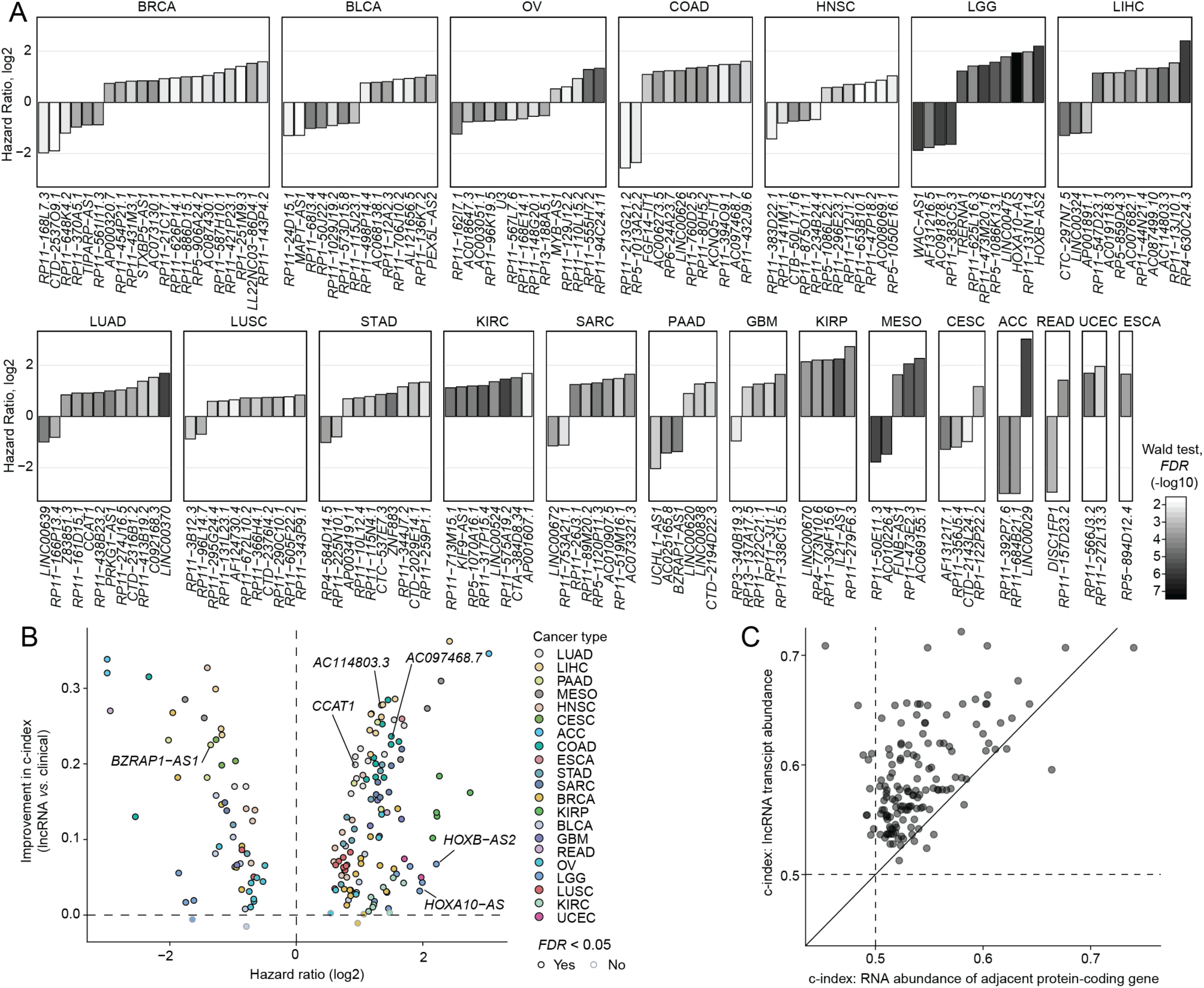
Elastic net proportional-hazards framework identifies 179 prognostic IncRNAs. **A.** The catalogue of 179 prognostic IncRNAs detected in 21 cancer types. IncRNAs are ordered by hazard ratios (HR) from the most to the least favourable in each cancer type and colored by statistical significance (Wald test, *FDR* < 0.05). **B.** Univariate prognostic models of 179 IncRNAs outperform baseline models of clinical variables in cross-validation experiments. Prognostic model performance is quantified using the concordance index (c-index). **C.** 179 IncRNAs show superior prognostic performance compared to adjacent protein-coding genes.

**Figure 3.**
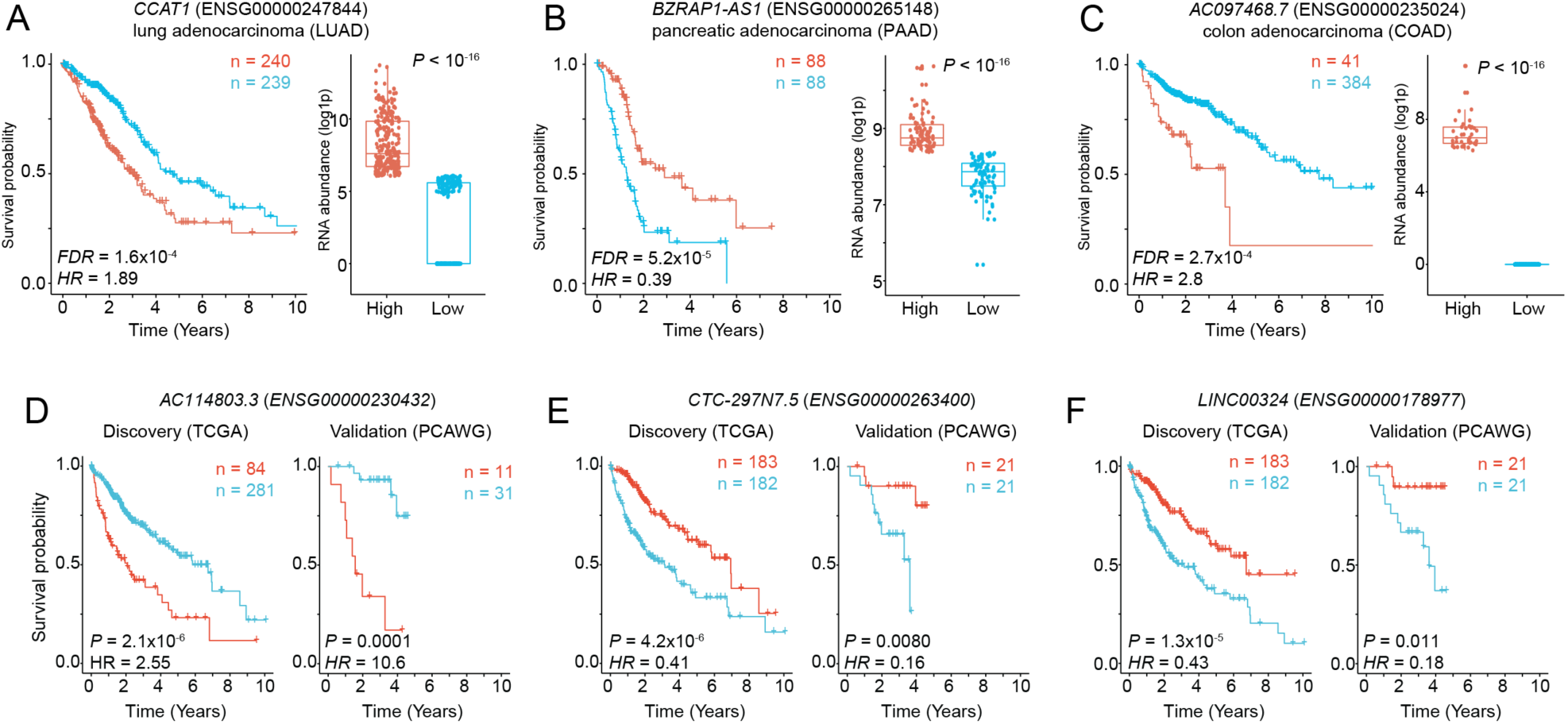
Examples of prognostic IncRNAs in multiple cancer types. **A-C.** Prioritized prognostic IncRNAs with Kaplan-Meier survival plots (left) and RNA abundance profiles as boxplots (right). Median-dichotomized transcript abundance profiles (FPKM-UQ) are shown (above median, red; below median, blue). **D-F.** prognostic IncRNAs in liver hepatocellular carcinoma (LIHC) with validation in an external cohort. Kaplan-Meier plots for the discovery cohort (left; TCGA) and the validation cohort (right, PCAWG) are shown. Sizes of patient groups are indicated in color. Hazard ratios (HR) and P-values were computed using Wald tests with no adjustment for covariates.

We prioritized 179 high-confidence prognostic lncRNAs in 21 cancer types that were detected as strong predictors in at least 50% of cross-validated models following the feature selection step of the elastic net framework (**Figure 2A, Supplementary Table 3**). The majority of lncRNAs (123/179 or 69%) were detected as unfavorable markers with respect to high transcript abundance (median HR = 2.3) while 56 lncRNAs were detected as favorable (median HR = 0.48). The largest numbers of prognostic lncRNAs were detected in multiple common cancer types: breast (21), bladder (14), ovarian (14), colorectal (12) and head and neck cancer (12). Lower-grade glioma (12) showed the strongest lncRNA candidates in terms of statistical significance. To quantify the 179 lncRNAs as prognostic markers individually and in combination with commonly used clinical variables, we separately considered each lncRNA regarding its prognostic model fit and also model performance in cross-validation experiments. The vast majority of individual lncRNAs (173/179) showed significantly higher prognostic accuracy across 1,000 cross-validations compared to baseline models comprising common clinical variables, with median increase of 0.11 in concordance index (Wilcoxon rank-sum test, *FDR* < 0.05; **Figure 2B**). Thus, our catalogue of lncRNAs provides complementary prognostic information to common clinical variables in a diverse set of human cancers.

We verified that our observed prognostic signals were specific to lncRNAs and did not solely reflect the prognostic signals of adjacent protein-coding genes. We identified 147 protein-coding genes located within ±10 kbps of 96/179 lncRNAs, including 106 genes that were antisense to lncRNAs (**Supplementary Table 4**). Prognostic models of lncRNA transcript abundance profiles exhibited higher concordance measures overall, compared to matching prognostic models of protein-coding genes (Rank-sum test, *P* = 7.88×10^−22^; **Figure 2C**). lncRNA-based prognostic models showed higher concordance index values in 139/147 cases compared to similar models of adjacent protein-coding genes, with a median improvement of 0.05 in c-index (c=0.58 for lncRNAs vs c=0.53 for protein-coding genes). Thus, the catalogue of prognostic lncRNAs is not transcriptionally confounded by adjacent protein-coding genes and represents a distinct non-coding search space for prognostic biomarker discovery.

### Top prognostic lncRNAs in cancer types of unmet need

We studied the 179 lncRNAs and the adjacent protein-coding genes for known associations with cancer. For example, *CCAT1* (ENSG00000247844) located in the chr8p24 super-enhancer locus is known to regulate *MYC* transcription through chromatin long-range interactions [34]. We found *CCAT1* as a marker of poor prognosis in lung adenocarcinoma (LUAD) (HR = 1.9, HR range = 1.4-2.5, Cox PH *FDR* = 1.6×10^−4^; **Figure 3A**). Overall, the 149 protein-coding genes located within ±10 kbps of the 179 lncRNAs included 10 known cancer genes of the Cancer Gene Census database [35] (*BCL10, HEY1, HOXA11, HOXA9, IRS4, LASP1, MYB, NCKIPSD, RNF43, SETD2*; Fisher’s exact *P* = 0.050), suggesting that a subset of the prognostic lncRNAs may be involved in the regulation of cancer driver genes through transcription regulatory and chromatin architectural interactions. Improved lncRNA-based survival predictions were found in several cancer types with poor outcomes that currently lack reliable prognostic biomarkers, such as colon, pancreatic and liver cancer. We reviewed the top candidates in these cancer types.

*BZRAP1-AS1* was found as a top significant lncRNA in the pancreatic adenocarcinoma cohort (PAAD) (ENSG00000265148; also known as *TSPOAP1-AS1*). Increased RNA abundance of *BZRAP1-AS1* associated with improved patient prognosis (HR = 0.39, HR range = 0.23-0.57, Cox PH *FDR* = 5.2×10^−5^; **Figure 3B**). Interestingly, *BZRAP1-AS1* is partially co-located in the genome with *RNF43*, a known driver gene with frequent mutations in pancreatic cancer (7%) and a potential therapeutic target [36, 37]. *RNF43* mRNA abundance alone did not appear prognostic in our dataset, potentially highlighting an independent function of this lncRNA. *BZRAP1-AS1* was recently reported as a survival-associated lncRNA in pancreatic cancer using a complementary transcriptomics dataset [38], validating our results obtained from the TCGA dataset.

*AC097468.7* was identified as a top significant lncRNA in the colon adenocarcinoma (COAD) cohort for its unfavorable transcript abundance profile. High abundance of *AC097468.7* (ENSG00000235024) in a minority of tumors (41/425 or 9.6%; median 1077 FPKM-UQ) was associated with worse prognosis (HR = 2.8, HR range = 1.7-4.9, Cox PH *FDR* = 2.7×10^−4^; **Figure 3C**), while the majority of tumors in the COAD cohort showed zero transcript abundance of the lncRNA and relatively better prognosis. The intergenic lncRNA is located between the genes *NHEJ1* and *IHH* within 10 kbps of both genes. *NHEJ1* is a core component of the non-homologous end joining (NHEJ) pathway that conducts DNA double strand break repair and maintains genome stability [39, 40]. Indian hedgehog (IHH) signaling regulates differentiation of colonocytes while epigenetic activation of IHH causes decreased self-renewal of colorectal cancer-initiating cells and increased sensitivity to chemotherapy [41][42]. We speculate that the prognostic lncRNA *AC097468.7* is involved in the regulation of these pathways through interactions with adjacent protein-coding genes. In summary, these examples demonstrate the potential of our catalogue to develop novel biomarkers and find functional lncRNAs for multiple important cancer types.

### Computational validation of *AC114803.3, CTC-297N7.5 and LINC00324* as prognostic lncRNAs in liver hepatocellular carcinoma

To investigate the 12 prognostic lncRNAs in hepatocellular carcinoma of the liver (LIHC), we studied an additional cohort of 42 patient tumors. The validation cohort was derived from the ICGC/TCGA Pan-Cancer Analysis of Whole Genomes (PCAWG) project [20] and was filtered to exclude tumors from TCGA. We found three lncRNAs with matching prognostic scores and significant P-values in both cohorts based on median dichotomization of transcript abundance values (*AC114803.3, CTC-297N7.5, LINC00324*) (**Figure 3D-F**).

*AC114803.3* was identified as a top significant lncRNA in both the discovery and the validation cohorts of liver cancer. Increased transcript abundance of this lncRNA was associated with worse prognosis in the TCGA cohort (HR = 2.6, HR range = 1.7-3.7, Cox PH *FDR* = 1.8×10^−5^) and confirmed in the PCAWG validation cohort (HR = 10.6, HR range = 3.1-36, *P* = 0.0001) (**Figure 3D**). In the discovery cohort, *AC114803.3* (*ENSG00000230432*) showed a discrete activation pattern with high transcript abundance in a minority of patients with poor prognosis (84/365 or 23% patient tumors with median 4239 FPKM-UQ) whereas a lack of RNA expression was observed in the other lower-risk group representing the majority of patients (0 FPKM-UQ). The discrete activation pattern was also observed in the validation cohort (11/42 tumors with median 0.093 FPKM-UQ, zero otherwise). *AC114803.3* is an antisense lncRNA co-located with the *PTPRN* gene that encodes a signaling protein and autoantigen in insulin-dependent diabetes [43]. A previous study found that DNA hypermethylation of *PTPRN* was associated with increased progression-free survival in ovarian cancer [44]. DNA hypermethylation is a repressive epigenetic mark inversely correlated with transcription, thus the study provides complementary evidence to our observation of high transcript abundance of the antisense lncRNA *AC114803.3* as a hazardous prognostic marker.

Two lncRNAs *CTC-297N7.5* and *LINC00324* were also found as markers of improved prognosis of LIHC through validation in the external dataset. Increased transcript abundance of *CTC-297N7.5* (*ENSG00000263400*) was associated with improved prognosis in the TCGA cohort (HR = 0.41, HR range = 0.29-0.61, *FDR* = 3.2×10^−5^) and validated in the PCAWG cohort (HR = 0.16, HR range = 0.032-0.76, *P* = 0.0080) (**Figure 3E**). *CTC-297N7.5* (also known as *TMEM220-AS1*) is an antisense lncRNA co-located with *TMEM220* encoding a poorly characterized transmembrane protein. This lncRNA has been reported recently as a prognostic factor in hepatocellular carcinoma [45], further validating our analysis. As the third prognostic lncRNA, increased transcript abundance of *LINC00324* (*ENSG00000178977*) was associated with improved prognosis in the TCGA LIHC cohort (HR = 0.43, HR range = 0.29-0.62, *FDR* = 6.0×10^−5^) and validated in the PCAWG cohort (HR = 0.18, HR range = 0.04-0.84, *P* = 0.011) (**Figure 3F**). This intergenic lncRNA has been functionally associated with the proliferation of gastric cancer cells [46]. Our computations validation analysis is limited by the available datasets and an overall lower detection of lncRNA transcript abundance in the PCAWG dataset. In summary, computational validation of our candidate lncRNAs in additional transcriptomics datasets and independent studies provides further support to these non-coding transcripts as prognostic biomarkers.

### Transcript abundance information of lncRNAs improves prognostic performance of known molecular and clinical tumor features

We asked whether the prognostic lncRNAs represented the transcriptomic footprints of well-defined clinical and molecular tumor subtypes. We investigated the statistical interactions of prognostic lncRNAs and of various molecular and clinical tumor annotations defined by TCGA [47]. We limited the analysis to a subset of lncRNAs (113/179) that were detected in 12/21 cancer types for which annotations of tumor features or subtypes were available in TCGA. We found 224 instances where transcript abundance of lncRNAs (36/113) associated with clinical or molecular tumor features (Chi-square test or Spearman correlation test, *FDR* < 0.05; **Figure 4A, Supplementary Table 5**). As expected, the majority of these features were also prognostic individually in univariate survival analyses (175/224 or 78%, Wald test, *FDR* < 0.05). The prognostic lncRNAs we identified in lower-grade glioma (LGG) associated with the largest number of molecular and clinical features, likely owing to well-defined subtypes of this form of brain cancer.

For example, transcript abundance profiles of the majority of prognostic lncRNAs in LGG were significantly associated with documented prognostic features such as *IDH* mutation status and *MGMT* promoter methylation [48, 49]. These data indicate that transcript abundance profiles of prognostic lncRNAs capture the transcriptomic signatures of known clinical subtypes and molecular features, further supporting the utility of these lncRNAs as prognostic biomarkers.

**Figure 4.**
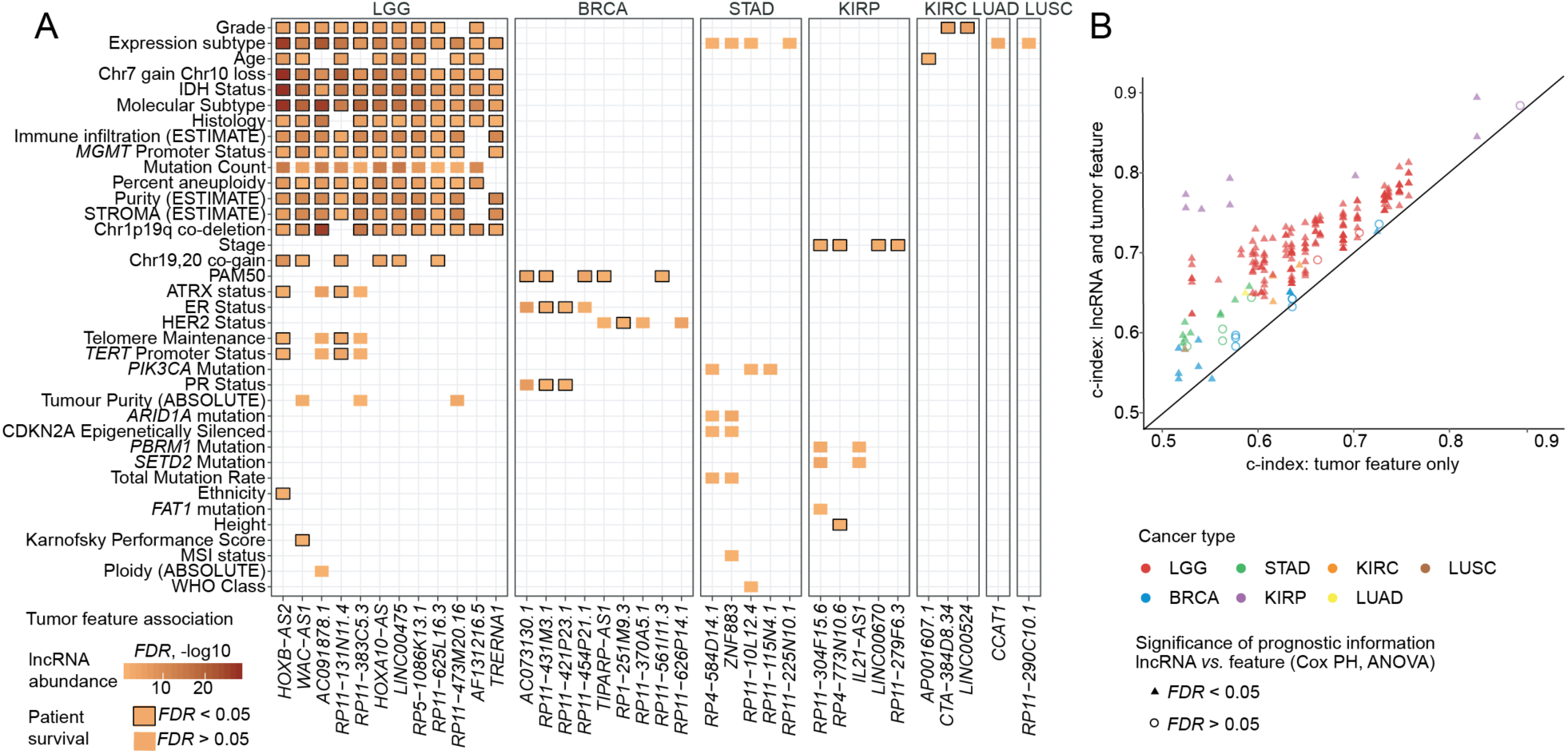
IncRNA transcript abundance improves prognostic performance of known molecular and clinical tumor features. **A.** RNA abundance of prognostic IncRNAs is associated with molecular and clinical tumor features and subtypes. Coloured boxes indicate significant associations with IncRNA transcript abundance (Chi-square test, *FDR* < 0.05). A subset of identified tumor features are also independently associated with patient survival (boxes with black frames; Wald test, *FDR* < 0.05). **B.** Combined prognostic models with IncRNA transcipt abundance and tumor features (y-axis) show consistently higher concordance values compared to baseline models with only tumor features (x-axis). Combined models with statistically significant contribution from IncRNA transcript abundance are indicated with triangles (Cox PH ANOVA, *FDR* < 0.05). Diagonal shows matching c-index values.

We asked whether the lncRNA transcript abundance profiles provided complementary information to clinical and molecular tumor features. We investigated the 224 cases where the 36 lncRNA transcript abundance profiles significantly associated with various tumor features, by comparing combined prognostic models (*i.e.*, lncRNAs and tumor features as predictors) with control prognostic models (*i.e.*, only tumor features as predictors) (**Figure 4B**). The majority of combined models (209/224 or 93%) showed improved prognostic performance and model fit (Cox PH ANOVA, *FDR* < 0.05). For example, combining the transcript abundance of lncRNA *RP11-279F6.3* (ENSG00000259641) with tumor stage resulted in an improved prognostic model in renal papillary cell carcinoma compared to a baseline model that only incorporated clinical stage as a predictor (median c = 0.93 *vs.* c = 0.87; Cox PH ANOVA *FDR* = 2.0×10^−4^). Similarly, transcript abundance of *RP5-1086K13.1* (ENSG00000224950) combined with co-deletion of chr1p and chr19q was a significantly better prognostic model in LGG compared to a baseline model that only used these chromosomal alterations for prediction (median c = 0.71 *vs.* c = 0.59, Cox PH ANOVA; *FDR* = 4.4×10^−7^). These results are limited by the molecular and clinical tumor features annotated by TCGA, as well as the lower overall transcript abundance and high tissue specificity of lncRNA transcription. Our analysis shows that integrating transcriptomic profiles of lncRNAs can improve the prognostic potential of previously established tumor features such as molecular subtypes and common genomic mutations.

### Prognostic lncRNAs in gliomas are associated with developmental, immune response and neurotransmission pathways

To study potential functional associations, we asked whether transcript abundance profiles of prognostic lncRNAs were associated with transcriptome-wide changes in high-risk tumors. For each lncRNA, we identified differentially regulated genes and mapped their biological context using pathway enrichment analysis [50]. The majority of prognostic lncRNAs (121/179 or 68%) associated with clear transcriptional signatures in lncRNA-stratified high-risk tumors, including at least 30 protein-coding genes with a two-fold change in transcript abundance (*FDR* < 0.05; **Supplementary Figure 4, Supplementary Table 6**). These genes were enriched in 3,048 GO biological processes and Reactome pathways in total (*FDR* < 0.01 from g:Profiler; **Supplementary Table 7**). The majority of detected pathways (75%) were enriched in the transcriptional signatures of a few lncRNAs (one to five) while a small subset of processes (5%) related to extra-cellular matrix organization were enriched in the signatures of more than 15 lncRNAs. This pancancer pathway analysis highlights the extent of functional diversity of high-risk tumors stratified by lncRNA abundance.

We studied the 12 prognostic lncRNAs identified in lower-grade glioma and evaluated their transcriptome-wide associations. We used a stringent approach that systematically accounted for the tumor mutation status of *IDH1/2* genes, a known marker of improved prognosis in glioma [51]. All groups of lncRNA-stratified high-risk LGG tumors were characterized by transcriptomic differences that were significant beyond *IDH* mutations (**Figure 5A**). To find pathways and processes commonly deregulated in these high-risk tumors, we performed an integrative analysis of the 12 lncRNA-stratified mRNA abundance signatures. This analysis revealed 325 biological processes and pathways that mapped to 1,345 protein-coding genes co-expressed with one or more of the 12 prognostic lncRNAs (ActivePathways [52] *FWER* < 0.05; **Figure 5B**). The pathway analysis highlighted 70 known cancer genes that were more frequently differentially expressed than expected from chance alone (Fisher’s exact test, *P* = 0.006; including key oncogenes *EGFR* and *TERT*). The pathway analysis revealed three broad functional themes: developmental processes (*e.g.*, forebrain development), immune system (*e.g.*, T-cell activation) and neurotransmitters (*e.g.*, trans-synaptic signaling). The majority of pathways (192/325 or 59%) were deregulated in the transcriptomic signatures of multiple prognostic lncRNAs, however only few pathways were apparent in all lncRNA-based transcriptomic signatures. These prognostic lncRNAs of LGG are co-regulated with diverse processes involved in brain development, neurotransmitter activity and tumorigenesis, suggesting that a subset of lncRNAs modulate cancer-related biological processes in brain tumors.

**Figure 5.**
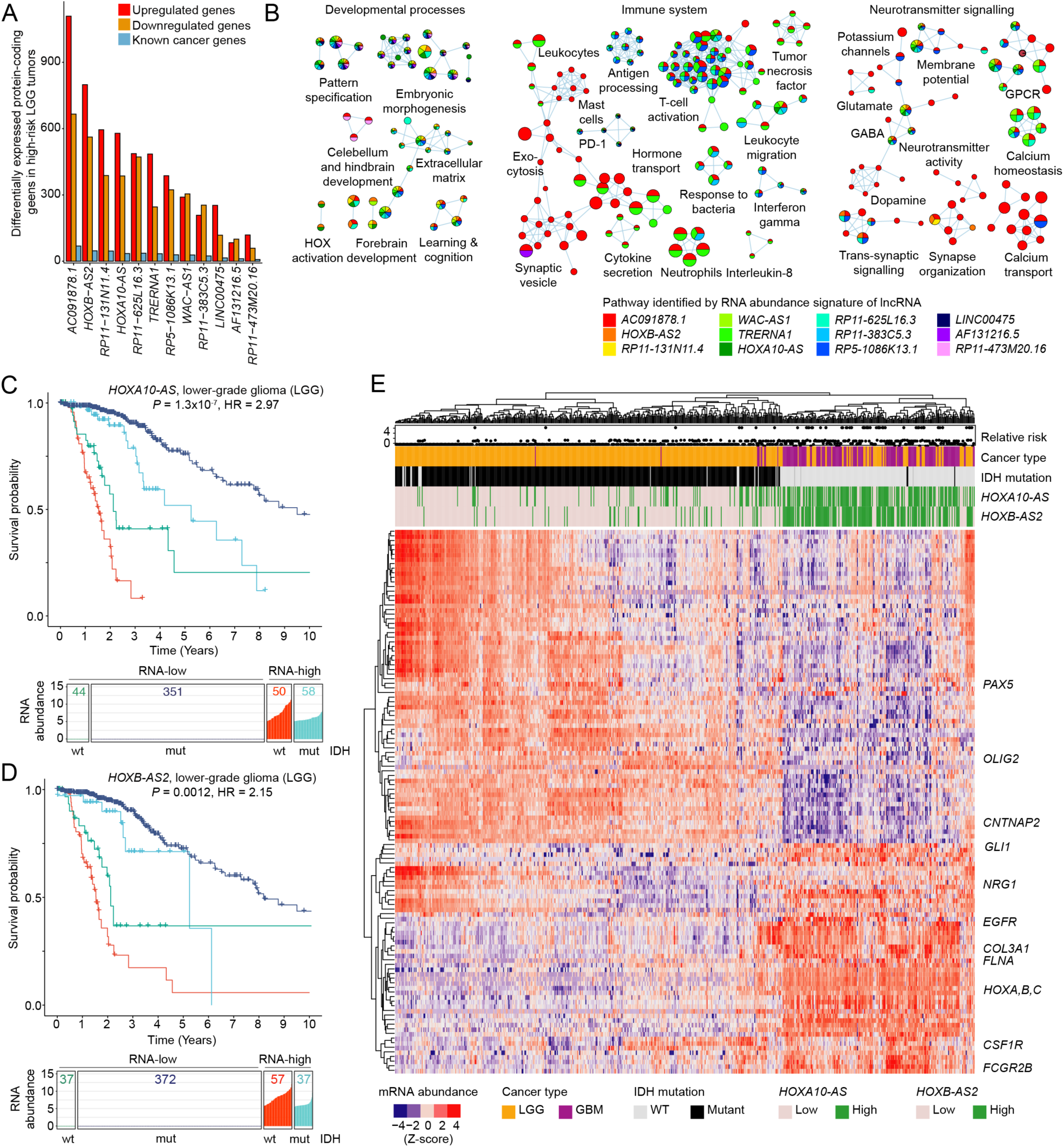
Prognostic IncRNAs in gliomas associate with deregulated driver genes and neurodevelopmental pathways. **A.** Prognostic IncRNAs in lower-grade glioma (LGG) associate with differential transcript abundance of protein-coding genes (*FDR* < 0.05), including many known driver genes. *IDH1/2* mutation status was modeled as a covariate in transcript abundance analysis. **B.** Pathway enrichment analysis of IncRNA-associated protein-coding genes in LGG shows de-regulation of neurodevelopmental, immune system and neuro-transmitter processes (ActivePathways *FWER*< 0.05). Enrichment map shows nodes as significantly enriched GO biological processes or Reactome pathways, nodes sharing many genes are connected with edges, and nodes are grouped by overall functional themes. Node colors represent the prognostic IncRNAs whose transcriptional signatures associated with the specific pathways. **D-E.** Transcript abundance of *HOXAIO-AS* and *HOXB-AS2* combined with *IDH1/2* mutation status improves prognostic models in LGG. Kaplan-Meier plots for *HOXAIO-AS* and *HOXB-AS2* (top) and distibutions of patients by transcript abundance (high *vs.* low; log 1 p FPKM-UQ) and *IDH* mutation status (wildtype *vs.* mutant) (bottom). Numbers indicate patient counts. **E.** *HOXA10-AS* and *HOXB-AS2* transcript abundance profiles define a malignancy gradient across LGG and glioblastoma (GBM). Heatmap shows differentially expressed genes in IncRNA-associated brain development pathways. High-risk LGGs with activated transcription of *HOXA10-AS* and *HOXB-AS2* cluster with GBMs and primarily include /DH-wildtype tumors. Known driver genes are shown.

### Transcript abundance of *HOXA10-AS and HOXB-AS2* defines a malignancy gradient across low- and high-grade gliomas

To further study the functional roles of prognostic lncRNAs in LGGs, we performed a transcriptome-wide comparison of lower-grade glioma and high-grade glioblastoma (GBM) tumors in TCGA. We focused on neurodevelopmental processes deregulated in high-risk LGG tumors apparent in our pathway analysis, such as the Reactome pathway *activation of anterior HOX genes in hindbrain development during early embryogenesis* that was enriched in mRNA signatures of high-risk tumors (*FWER* = 0.003). In this pathway, developmental transcription factors *HOXA1, HOXA2, HOXA3, HOXA4* and *HOXC4* were co-activated with the two prognostic lncRNAs *HOXA10-AS* and *HOXB-AS2* in high-risk LGG tumors. An extended set of significantly enriched GO processes related to brain and central nervous system development was also found. These processes included 118 differentially expressed genes including known brain cancer genes *EGFR, GLI1*, and *CNTNAP2*. The potential neurodevelopmental mechanisms altered in high-risk gliomas highlighted the HOX-associated lncRNAs as high-priority targets for further study.

*HOXA10-AS* transcript abundance appeared as highly hazardous in the LGG cohort (HR = 3.8, HR range = 2.38-5.19, Cox PH *FDR* = 5.0×10^−8^) and a similar highly significant association was observed for *HOXB-AS2 (*HR = 4.6, HR range = 2.2−5.1, FDR = 1.4×10^−6^). When combined with *IDH* mutation status, zero-dichotomized transcript abundance profiles of *HOXA10-AS* and *HOXB-AS2* improved LGG prognostic models compared to univariate models with *IDH* mutation status alone (HR = 2.97, HR range = 2.0-4.4, *FDR* = 8.0×10^−7^, and HR = 2.15, HR range = 1.4-3.4, *FDR* = 0.002, respectively) (**Figure 5C-D**). In particular, the subset of ∼10% LGG patients with no *IDH* mutations and high lncRNA abundance were stratified as the highest-risk group compared to all other patients. Thus, the two lncRNAs may represent novel molecular biomarkers of advanced LGGs whose discrete transcriptional activation patterns in combination with *IDH* mutation status indicate dismal outcome.

We quantified the transcriptional activation of *HOXA10-AS* and *HOXB-AS2* in lower-grade gliomas and glioblastomas. Hierarchical transcriptome clustering of *HOXA10-AS* and *HOXB-AS2* together with the 118 developmental genes across the LGG and GBM cohorts revealed a malignancy gradient of gliomas (**Figure 5E**). The major low-risk cluster of tumor transcriptomes contained LGGs with little or no transcription of the two prognostic lncRNAs. In contrast, the cluster of high-risk LGGs was clearly defined by an increased abundance of *HOXB-AS2* and *HOXA10-AS*. This high-risk set of LGGs was clustered together with GBMs, while GBMs were defined by even higher transcript abundance of the two prognostic lncRNAs as well as oncogenes such as *EGFR* and *GLI1*. In LGG, *HOXB-AS2* and *HOXA10-AS* were characterized by bimodal transcript abundance: high transcript abundance was observed in few tumors (19% and 21% respectively), and silencing with zero transcript abundance of the two lncRNAs in the majority of tumors. Further, the majority of GBM tumors showed high transcript abundance of *HOXB-AS2* (68%) and *HOXA10-AS* (70%) and their overall transcript abundance was higher in GBMs than in LGGs (**Supplementary Figure 5**), indicating that HOX-antisense lncRNA expression positively correlated with tumor grade. *HOXB-AS2* and *HOXA10-AS* were not significant prognostic in the GBM cohort, perhaps owing to the overall poor prognosis of these advanced tumors (**Supplementary Figure 6**). This neurodevelopmental gene signature may represent a transcriptomic subtype of LGG that is marked by discrete transcriptional activation of the two HOX-antisense lncRNAs with prognostic relevance and functional roles.

**Figure 6.**
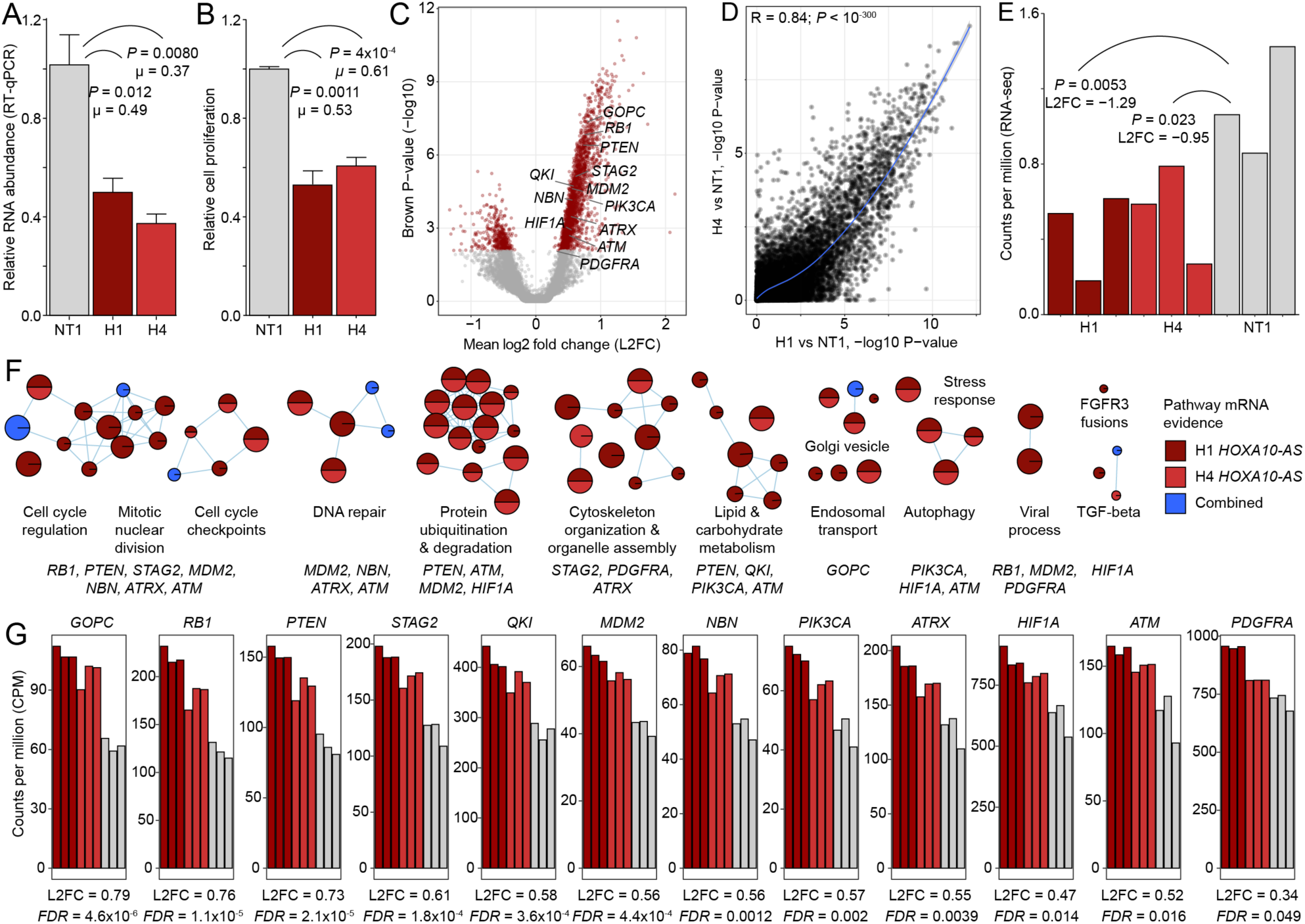
*HOXAIO-AS* knockdown in patient-derived GBM caused reduced cell proliferation and deregulation of cell cycle genes and glioma drivers. **A.** siRNA knockdown of *HOXA10-AS* caused its reduced transcript abundance, as shown by RT-qPCR. Knockdown was performed in triplicates with two siRNAs (H1, H4) targeting the IncRNA and a non-targeting control siRNA (NT1). Significance (Welch T-test) and normalized mean values μ are shown. **B.** *HOXA10-AS* knockdown caused reduced cell proliferation on day six post-transfection. C. Down-regulation of *HOX10A-AS* in siRNA experiments was confirmed using RNA-seq. **D.** Transcriptome-wide changes induced by the two siRNAs (H1, H4) were strongly correlated. Pearson correlation and loess trendline are shown. **E.** Volcano plot shows protein-coding genes with significant mRNA abundance changes in *HOX1*0*A*-*AS*-inhibited cells. High-confidence glioma genes are highlighted. **F.** *HOX10A-AS* -inhibited cells showed deregulation of biological processes (ActivePathways *FWER* < 0.05). Enrichment map shows enriched pathways as a network with nodes as pathways and edges connecting pathways with many shared genes. Node color indicates the siRNA experiment that led to differential expression of the pathway. **G.** *HOX10A-AS* inhibition caused transcriptional activation of glioma driver genes. FDR-adjusted Brown P-values and mean log2 fold-change values (L2FC) are shown.

### *HOXA10-AS* knockdown in patient-derived glioblastoma cells reduces proliferation and deregulates cell cycle genes and glioma drivers

The prognostic and pathway associations of *HOXA10-AS* transcript abundance prompted us to investigate this lncRNA functionally. We performed a siRNA-mediated knockdown experiment of *HOXA10-AS* followed by a six-day cell proliferation experiment using the primary patient-derived GBM cell line G797 [53, 54]. To minimize off-target effects on the protein-coding gene *HOXA10* antisense to the lncRNA, we used two siRNAs against the unique exon three of the lncRNA. siRNA-mediated inhibition of *HOXA10-AS* led to two-fold reduction in transcript abundance of the lncRNA relative to non-targeted controls (T-test, *P* ≤ 0.020; **Figure 6A**). *HOXA10-AS* -inhibited cells showed ∼40% lower cell proliferation at the 6-day timepoint (*P* ≤ 0.0011; **Figure 6B**). Transcriptional inhibition of *HOXA10-AS* and the resulting reduction in cell proliferation was robustly observed in experiments conducted with either of the two targeting siRNAs. These findings indicate the function of *HOXA10-AS* in regulating cell proliferation in glioma and confirm a recent report on this lncRNA [55].

To further understand the role of *HOXA10-AS* in the hallmark pathways of glioma, we conducted whole-transcriptome RNA sequencing (RNA-seq) of *HOXA10-AS* depleted cells three days after siRNA transfection. We found a pronounced transcriptional response of 2,428 differentially expressed genes in *HOXA10-AS*-inhibited cells relative to non-targeted controls (Brown *FDR* < 0.05, log2 fold-change > 1.2 using TREAT [56]; **Figure 6C**). The two targeting siRNA induced highly correlated transcriptome-wide changes (Pearson correlation test, *R* = 0.84, *P* < 10^−300^) and confirmed reduced transcript abundance of *HOXA10-AS* in siRNA-treated cells (*P* < 0.023, L2FC < −0.95; **Figure 6D-E**). We interpreted the transcriptomic changes induced by *HOXA10-AS* knockdown using pathway enrichment analysis and found 84 biological processes and molecular pathways enriched in the differentially expressed genes (*FWER* < 0.05 from ActivePathways [52]; **Figure 6F**). The pathways and processes were associated with 2,108 differentially expressed genes through the sensitive data fusion approach implemented in ActivePathways. Known cancer genes were significantly enriched (137 observed vs 91 expected, Fisher’s exact *P* = 2.4×.10^−7^) and included 12 up-regulated genes that are well recognized in the biology and mutational driver landscape of glioma (*GOPC, RB1, PTEN, STAG2, QKI, MDM2, NBN, PIK3CA, ATRX, HIF1A, ATM, PDGFRA*; **Figure 6G**) [35, 57, 58]. For example, the enriched GO process *regulation of mitotic cell cycle* (*FWER* = 0.03) provides an explanation to our observed phenotype of reduced glioma cell proliferation and implicates *HOXA10-AS* in the transcriptional rewiring of cell proliferation pathways. 128 genes of this pathway were deregulated in *HOXA10-AS* inhibited cells, including two upregulated tumor suppressors *RB1* and *PTEN*. Additional enriched pathway themes such as DNA repair (*MDM2, NBN, ATRX, ATM*), protein ubiquitination (*PTEN, ATM, MDM2, HIF1A*), lipid metabolism (*PTEN, QKI, PIK3CA, ATM*) and TGF-beta signaling (*HIF1A*) suggest further roles of *HOXA10-AS* in mediating cell proliferation in glioma. Finally, we asked whether our observed transcriptional and proliferative differences of *HOXA10-AS* depleted cells would be explained by the antisense homeobox gene *HOXA10* that modulates the tumorigenic potential of glioblastoma stem cells [59]. *HOXA10* showed no significant differences in transcript abundance in *HOXA10-AS* depleted cells compared to control-transfected cells in RNA-seq data and RT-qPCR assays (**Supplementary Figure 7**), suggesting that our functional and transcriptional evidence of altered cell proliferation is specific to the lncRNA *HOXA10-AS* and is not significantly confounded by any off-target effects of our knock-down experiment. In summary, these findings provide functional evidence to one of our predicted prognostic lncRNAs as a regulator of hallmark cancer processes in glioma.

## DISCUSSION

The current knowledge of cancer driver genes and molecular classifiers is primarily derived from the protein-coding genome while the vast non-coding genome remains understudied. Our findings of lncRNAs as prognostic factors in multiple cancer types are consistent with the increasing appreciation of lncRNAs in diverse cellular processes and human diseases. Our study highlights a facet of the non-coding genome that has great potential for basic and translational discoveries. Our machine learning analysis identified a subset of lncRNAs as robust predictors of patient survival in cross-validation experiments, suggesting that these transcripts should be further evaluated as prognostic biomarkers in diverse molecular datasets. To establish one lncRNA as a *bona fide* modulator of cancer hallmark processes, we functionally validated a prominent candidate lncRNA *HOXA10-AS* in patient-derived glioblastoma cells and observed significantly reduced cell viability upon lncRNA depletion, differential expression of glioma driver genes as well as transcriptome-wide changes enriched in proliferative, DNA damage response and metabolic pathways. These data suggest further functional and mechanistic experiments to validate *HOXA10-AS* as a potential therapeutic target. The integrative analysis and experimental validation data lend confidence to our overall catalogue of lncRNAs. However, our analysis remains inconclusive to whether all or most candidate lncRNAs are functional in cancer cells or alternatively represent passive indicators of transcriptional activity. On the one hand, functionally inactive ‘passenger’ lncRNAs may be modulated transcriptionally or epigenetically as part of global gene regulatory programs that control hallmark cancer pathways such as proliferation. These markers of large regulatory programs would be expected to outperform any prognostic models based on individual protein-coding genes. For example, we observed that a subset of lncRNAs with hazardous risk profiles were sharply up-regulated in high-risk tumors and completely silenced in lower-risk tumors. These lncRNAs may be epigenetically repressed in the majority of tumors and aberrantly activated in the high-risk minority group of tumors. Such a binary zero-dichotomization pattern is a promising property for biomarker development owing to a natural threshold separating high-risk and low-risk patients, although further validation in independent cohorts is required. On the other hand, a subset prognostic lncRNAs may be functional in cells and act as functional ‘drivers’ that activate oncogenic processes or inhibit tumor suppressive pathways through interactions with DNA, RNA and proteins. However, further experiments are needed to validate the prognostic lncRNAs as drivers of cancer phenotypes, such as large-scale genome editing screens that are increasingly targeting the non-coding genome encoding lncRNAs [60]. Our findings of prognostic lncRNAs are ultimately limited by the transcriptional and clinical information that was available for inference and validation. The TCGA tumor cohorts that we studied are under-represented in rare and early-stage malignancies and the available clinical variables and patient follow-up data are limited. It is plausible that lncRNA transcription in cancers is associated with unrecorded environmental, genetic and phenotypic variables that confounded our inference of prognostic markers. We used RNA-seq datasets that had been optimized for mRNA quantification and thus additional lncRNAs likely remain uncharacterized or lie below the detection limit of RNA-sequencing protocols. Future multi-omics datasets with deep clinical profiles of patients will enable further discoveries and validation of non-coding RNAs. Our study is a step towards systematic characterization of non-coding RNA genes as molecular biomarkers and functional regulators of oncogenesis.

## ACKNOWLEDGMENTS

We would like to thank Dr. Fritz Roth, Dr. Hansen He, Hassaan Maan and members of the Reimand lab for thoughtful discussions. This research was partially funded by the Natural Sciences and Engineering Research Council of Canada (NSERC) Discovery Grant to J.R., MBP Excellence Award to K.I. from the University of Toronto Department of Medical Biophysics, and the Ontario Institute for Cancer Research (OICR) Investigator Award to J.R, and the OICR Brain Tumour Translational Research Initiative to J.R., D.S., and P.B.D. Funding from the OICR is provided by the Government of Ontario. The results published here are in part based upon data generated by the TCGA Research Network: https://www.cancer.gov/tcga.

## AUTHOR CONTRIBUTIONS

K.I. led computational analyses and developed the methodology. K.I. and C.L. analyzed the data. K.I., L.J., D.S., and J.R. interpreted the data. L.J. and R.T. conducted experiments. D.S. supervised the experiments. K.I. and J.R. conceived and designed the study. F.C. and P.B.D. contributed patient-derived cell lines and know-how. J.R. supervised the study. K.I. and J.R. wrote the manuscript with input from all authors. All authors approved the final manuscript.

## METHODS

### Data Collection

We downloaded RNA-seq data of the TCGA project for 32 tumor types from the Genome Data Commons (https://portal.gdc.cancer.gov). Overall survival data was retrieved from the latest publication of the TCGA PanCanAtlas project [22, 29]. We selected 29 cancer types where cohorts of at least 50 patients were available. We only analyzed one tumor specimen per patient and maintained the tumor with a smaller TCGA serial number for patients with multiple specimens. Additional information on patient clinical variables such as alcohol consumption, smoking status and molecular subtypes was downloaded using the R package TCGABiolinks [47]. We intersected clinical information and transcript abundance data for each cancer type and retained patient cohorts where matched datasets were available. For lncRNA annotations, we downloaded the latest comprehensive annotation set of 5’ lncRNA CAGE peaks from the FANTOM-CAT project [2]. We studied 5,785 lncRNAs that were annotated by FANTOM-CAT and the ENSEMBL database and for which RNA abundance data were available in TCGA.

### Processing TCGA RNA-seq data

For all cancer types of the TCGA dataset, we retrieved processed RNA-seq files as FPKM-UQ measurements and raw counts from the Genome Data Commons website. lncRNAs often have low transcript abundance and we first removed the lncRNAs that were not detected in any patient tumor sample across all cohorts in TCGA RNA-seq data (n=94). Further, we evaluated median transcript abundance of each lncRNA in every cancer type and included two classes of lncRNAs in further analyses. First, we included lncRNAs with a median FPKM-UQ above 0. Second, we also included a set of lncRNAs with binary transcript abundance profiles. These lncRNAs showed median transcript abundance of zero FPKM-UQ representing the majority of tumor samples, while a minority of tumor samples (at least 15) showed transcript abundance of at least 100 FPKM-UQ. To evaluate tissue specificity of lncRNA transcription profiles, we used the UMAP (Uniform Manifold Approximation and Projection) dimension reduction method [31] and the corresponding R package to perform clustering of log1p-transformed FPKM-UQ lncRNA transcript abundance values across the entire TCGA cohort.

### Training survival models and evaluating generalizability

For each cancer type, we evaluated the association between all lncRNAs and overall patient survival. We also evaluated the association between available clinical variables and overall survival for comparison. For each cancer type, we split samples randomly into two groups, with 70% as the training set and 30% as the test set. Patients within each training cohort were median-dichotomized by the transcript abundance of each lncRNA. In case of lncRNAs with median transcript abundance of zero, patients with lncRNA transcript abundance above zero were labeled as high-abundance and those with zero abundance were labeled as low-abundance. We used the elastic net framework with a Cox proportional hazards link function to train patient survival models and to perform feature selection. All univariate models were built using the R package “survival”. Elastic net modelling was performed using the R package “glmnet” where the penalty hyperparameter λ was determined by fivefold cross-validation within each training set. We used the fixed hyperparameter value α=0.5 for the elastic net model. We employed 1000-fold cross-validation with 70/30% random split of training and testing data for each cancer type. Within each fold, initial elastic-net multivariate models included as predictors all lncRNAs that were univariately survival-associated in the training set (univariate Cox proportional-hazards (PH) *P*<0.05). Feature selection during model fitting and regularization determined a non-redundant subset of lncRNAs as predictors in the training data. Subsequent cross-validation evaluated the models using concordance index (c-index), an accuracy measure extended to survival analysis [33]. The multivariate Cox PH elastic net models were then applied to the remaining 30% of the test set to obtain a concordance index (c-index) using the R package “survcomp”. Besides lncRNA-based predictors, clinical variables that were available for each cancer types were also used to build a multivariate model using the training set and applied on test set in a similar manner. Of clinical variables, patient age was always available for all tumor types in TCGA, while other features such as tumor stage, grade and ethnicity were available for a subset of cancer types. Lastly, the available clinical variables were integrated with the lncRNA transcript abundance profiles selected by the elastic net into one multivariate model (the combined model) that was also trained and tested separately. Thus, there were three distinct performance metrics (c-indices) obtained overall for each round of training. The entire outlined process was repeated 1000 times, randomly splitting the data at each iteration. For each cancer type, we subsequently compared the three distributions of c-indices using the two-sided U test to a set of reference models that only utilized clinical variables for survival predictions. Finally, to assess the performance of our models on random data, we shuffled survival outcome across all TCGA patients of a given cancer type while maintaining the order of all predictor variables (lncRNAs and clinical variables). This permutation strategy disrupted the association of survival information and molecular and clinical predictors, The analysis of this simulated data allowed us to evaluate the statistical calibration of our method. We generated 100 random datasets and conducted 100 cross-validations on each of these datasets. We compared c-indices between models fitted using shuffled outcome data and real outcome data using a two-sided U-test. As expected, we found considerably lower performance of our models on random data that centered on the expected performance values of random predictors (c≈0.5), indicating that our models were well calibrated and not prone to statistical inflation and overfitting.

### Selecting top prognostic lncRNAs

To prioritize lncRNAs, we summarized the number of times each lncRNA was maintained as a prognostic feature in all the elastic-net survival models across cross-validations. To obtain the most consistent candidates, we considered the lncRNAs in each cancer type that were included in at least 50% (≥500/1000) of iterations. This list of lncRNAs was further evaluated individually. For validation, we fitted multivariate Cox PH models using each lncRNA candidate together with available clinical variables in respective cancer cohorts to confirm that the prognostic effect of lncRNAs remained present even when accounting for common clinical variables. We also evaluated Schoenfeld residuals to confirm that the proportionality assumption of the Cox-PH model was met (**Supplementary Table 3**). Finally, we removed a small subset of candidate lncRNAs that showed opposing hazards in different cancer types. To evaluate the performance of individual lncRNA candidates within the TCGA dataset, we conducted a second round of internal cross-validation. Using one lncRNA candidate at a time, we split the respective cancer patient cohort into training (70%) and testing samples (30%) as described above. Univariate Cox PH models were fitted and evaluated on the test datasets to obtain a distribution of c-indices for each lncRNA candidate. Similarly, we conducted internal cross-validation of clinical variables as a baseline reference, by fitting multivariate Cox PH models and evaluating their performance on test sets using the c-index. We also compared combined models where clinical variables were used together with lncRNA transcript abundance profiles for patient survival prediction. These distributions of c-indices were compared using the two-sided Wilcoxon rank-sum tests and resulting P-values were adjusted using the Benjamini-Hochberg false discovery rate (FDR) procedure [61].

### Validating prognostic lncRNAs in additional cohort of hepatocellular carcinoma

We used an independent dataset of transcriptomics and patient clinical information available in the ICGC/TCGA Pan-cancer Analysis of Whole Genomes (PCAWG) project [20]. We focused on the liver cancer cohort and removed any patient samples profiled in the TCGA project to create an entirely independent validation cohort comprising primarily of liver cancers (hepatocellular carcinomas, HCC) of Japanese individuals [62], resulting in a cohort of 42 tumors with uniformly processed RNA-seq data [63]. Twelve lncRNAs identified in the TCGA LIHC cohort were queried for prognostic signals in the validation cohort. Within the validation cohort, we considered lncRNAs with FPKM-UQ values greater than 0.05 measured in at least five patients. We dichotomized patients by lncRNA transcript abundance as described above. To evaluate significance of patient survival associations, we fitted univariate Cox-PH models with binary predictors reflecting lncRNA transcript abundance and plotted their Kaplan-Meier survival curves using the ‘Survival’ and ‘survminer’ packages in R. We considered those lncRNAs with nominal P-values from Wald tests as significant (*P* < 0.05).

### Comparing survival associations of lncRNAs and adjacent protein-coding genes

We identified protein-coding genes that were located within 10,000 bps of lncRNA genes using the Genome Reference Consortium Human Build 38 (GRCh38) and the bedtools software [64]. We identified pairs of 96 lncRNAs and 147 protein-coding genes that we evaluated further for differences in patient survival associations. For each pair, we fitted univariate Cox-PH models using median-dichotomized lncRNA transcript abundance labels as described above, and compared these to Cox-PH models fitted using median-dichotomized transcript abundance values of corresponding protein-coding genes. We compared the sets of two models using cross-validation performance (i.e., c-indices) and also model fits (i.e., FDR-adjusted *P*-values from the Wald test). We also fitted multivariate models using transcript abundance values of both the protein-coding gene and the lncRNA gene, and compared those models to univariate models of protein-coding genes using ANOVA. Multiple testing correction was performed using the Benjamini-Hochberg FDR procedure.

### lncRNA associations with clinical and molecular tumor subtypes

We conducted a systematic analysis of clinical and molecular subtypes of TCGA tumors using data curated in the R package TCGABiolinks [47]. These clinical and molecular features included basic clinical variables included in our elastic net framework described above (patient age, sex and tumor stage and/or grade, *etc*. as available in TCGA), and additional variables such as molecular subtypes, specific prognostic mutations and tumor histology annotations. These comprehensive sample-specific annotations were only available for 12/21 cancer types for which high-confidence prognostic lncRNAs were predicted, and we further analyzed only the 113/179 lncRNAs predicted in these cancer types. For each lncRNA, we evaluated whether the transcript abundance was significantly associated with clinical and molecular features. Dichotomized lncRNA transcript abundance profiler (high *vs.* low) were compared to clinical and molecular features using chi-squared tests as most clinical and molecular variables per patient were recorded as binary categories. For numerical clinical and molecular variables (such as age), we analyzed the spearman correlation between the variables and lncRNA transcript abundance. We adjusted P-values for multiple testing using the Benjamini–Hochberg FDR procedure and selected significant associations (*FDR* < 0.05). All clinical features from the analysis that were significantly associated with our lncRNA candidates were also evaluated for associations with overall patient survival. For the lncRNAs associated with at least one clinical or molecular feature, we extracted the corresponding (c-index) from a Cox-PH model (model 1). Next, we fitted univariate Cox-PH models with the clinical or molecular feature as a predictor of overall patient survival within the respective cancer cohort. For each model we extracted its c-index, HR and Wald test *P*-value (model 2). Finally, we fitted a multivariate model with both the clinical or molecular feature with the lncRNA transcript abundance profile that it was associated with (model 3). This allowed us quantify the combination of lncRNA transcript abundance and previously annotated clinical and molecular features. Tests with Cox PH models were defined as:

<u>Test #1</u>: Anova (model 1, model 3), to assess the improvement of the survival association when using both lncRNA transcript abundance and clinical/molecular features as predictors, compared to lncRNA-based predictors alone.
<u>Test #2</u>: Anova (model 2, model 3), to assess the improvement of the survival association when using both lncRNA transcript abundance and clinical/molecular features as predictors, compared to clinical and molecular features as predictors alone.

To obtain the final list of lncRNA-associated clinical and molecular features that showed significant improvement in survival association in combination with lncRNA transcript abundance, we considered two criteria: a significant likelihood ratio test (*FDR* < 0.05) from the Test #2 above, and an absolute increase in c-index in cross-validation experiments.

### Pathway enrichment analysis of lncRNA-associated protein-coding genes

For each prognostic lncRNA, tumors of a given type were first classified as high-risk or low-risk, based on median dichotomization of the lncRNA as described above. We conducted differential transcript abundance analysis to identify protein-coding genes that were differentially expressed in high-risk tumors. We used raw sequencing read counts from the TCGA RNA-seq datasets and applied the Limma method for differential transcript abundance analysis [65]. We considered all protein-coding genes with a filter on effect size (absolute fold change (FC) > 2, *FDR* < 0.05). We highlighted known cancer genes curated in the COSMIC Cancer Gene Census dataset [35]. We then used g:Profiler web server [66] to identify significantly enriched Reactome pathways and GO biological processes in the differentially expressed protein-coding genes associated with each lncRNA. We filtered gene sets (pathways and processes) to only include at least 10 and less than 250 annotated genes and a minimum of five pathway-annotated genes differentially expressed in the lncRNA-stratified set of high-risk tumors. Pathway enrichments were filtered by statistical significance (*FDR* < 0.05 in g:Profiler). An additional stringent version of this analysis was conducted for the 12 prognostic lncRNAs in LGG. First, protein-coding genes with differential mRNA abundance were detected in the LGG cohort by specifically accounting for IDH mutation status as covariate in the Limma framework. Second, pathway enrichment analysis was conducted using the data fusion approach implemented in the ActivePathways package [52]. ActivePathways prioritized protein-coding genes that showed differential transcriopt abundance signals for multiple prognostic lncRNAs in the LGG cohort. All nominally significant genes were considered for input pathway enrichment analysis according to default parameter settings of ActivePathways (gene-based Brown *P*<0.1). Resulting enriched pathways were adjusted for multiple-testing correction and filtered according to default settings (ActivePathways, Holm family-wise error rate (*FWER)*<0.05). Pathway enrichment maps were built in Cytoscape using standard procedures and manually curated for groups of related pathways as functional themes [50]. For LGG, we focused on a subset of neurodevelopmental pathways and associated protein-coding genes for further enquiry into the top prognostic lncRNAs in LGG, *HOXA10-AS* and *HOXB-AS2.* We generated heatmaps to summarize the expression of these genes in LGG and GBM using the “ComplexHeatmap” package [67]. The heatmap was generated using log1p transformed FPKM-UQ values and a hierarchical clustering with Pearson correlation distance was applied. Relative risk was calculated for LGG patients using a multivariate Cox-PH model accounting for dichotomized transcript abundances of both *HOXB-AS2* and *HOXA10-AS*.

### Cell Culture of patient-derived GBM cell lines

The human glioma G797 cells were prepared as described previously as a bulk patient-derived cell cultures [53, 54]. We selected the G797 patient-derived cell line as a suitable candidate for our experiments based on previously generated RNA-seq data [53] that indicated a relatively high native transcript abundance of lncRNA *HOXA10-AS* in these cells. G797 cells were maintained in serum-free NeuroCult™ NS-A Basal Medium (STEMCELL Technologies Canada Inc) supplemented with N2, B27, EGF (10 ng/ml), and FGF-2 (10 ng/ml), as described previously [68].

### siRNA mediated knockdown of HOXA10-AS

A TriFECTa DsiRNA kit (hs.Ri.HOXA10-AS.13) containing one non-targeting control DsiRNA (NT1) and DsiRNAs targeting *HOXA10-AS*, and an additional DsiRNA targeting *HOXA10-AS* (CD.Ri.209973.13.8) were purchased from Integrated DNA Technologies. The targeting sequences were: H1, AGACGATTTCAACTGAAGTAATGAA; and H4, GGTACCTGGAGACGATTTCAACTGA. Transfection of DsiRNAs was performed using Lipofectamine RNAiMAX Reagent (Thermo Fisher Scientific) as per manufacturer’s protocol. Exon3 of *HOXA10-AS* is directly antisense to protein-coding exons of *HOXA10*. To avoid off-target effects of knocking down *HOXA10-AS*, we purposefully avoided this region in siRNA design and instead selected siRNAs targeting exon2 of *HOXA10-AS*, a region unique to *HOXA10-AS* and not overlapping with *HOXA10*. We confirmed successful knock-down of *HOXA10-AS* by RT-PCR using primers flanking exon2 of *HOXA10-AS*. With depletion of *HOXA10-AS* we did not observe a significant change in *HOXA10* transcript abundance in either our RT-qPCR or RNA-seq experiments.

### PrestoBlue Cell Viability assay

PrestoBlue Cell Viability assays (A13262, Thermo Fisher Scientific) were performed as per manufacturer’s protocol. Briefly, 5,000 cells were seeded into each well of 96-well plates on day 0 of DsiRNA transfection. On each day of viability assay, cells were incubated with 100ul fresh complete medium with the PrestoBlue reagent for 40 min. Then the fluorescence readout was obtained using a SpectraMax Gemini EM Microplate Reader (Molecular Devices) with the excitation/emission wavelengths set at 544/590 nm. Cell viability is reported at the 6-day timepoint of the experiment.

### RNA isolation, cDNA synthesis, and real-time QPCR analysis

RNA samples were extracted from cells three days post DsiRNA transfection using Quick-RNA Microprep Kit (Zymo Research), treated with DNase I (Zymo Research), quantified using the Qubit, and reverse transcribed into cDNA using SuperScript IV VILO (Invitrogen). Primers were designed to span and/or overlap exon junctions using Primer3Plus. Primers were validated against a standard curve and relative mRNA expression levels were calculated using the comparative Ct method normalized to PPIB mRNA [69]. Real-time quantitative PCR (qRT-PCR) reactions were performed on an CFX384 (Biorad) in 384-well plates containing 12.5 ng cDNA, 150 nM of each primer, and 5 μl of 2X SensiFAST SYBR No-ROX kit (Bioline) in a 10 μl total volume. The following RT-qPCR primers were used: *HOXA10-AS* (NR_046609.1; forward: CAGAGAGAAGGGTGGAGGTG; reverse: CTCAGGAGCCTCGTGTCTTT), *HOXA10* (NM_018951.3; forward: CCTTCCGAGAGCAGCAAAG; reverse: TGCGTTTTCACCTTTGGAAT), control gene *PPIB* (NM_000942.4; forward: GGAGATGGCACAGGAGGAA; reverse: GCCCGTAGTGCTTCAGTTT).

RNA-seq libraries were prepared using Illumina TruSeq Stranded mRNA Sample Prep Kit (20020594) as per manufacturer’s protocol. The barcoded cDNA libraries were then checked with Agilent Fragment Analyzer for fragment size and quantified with ddPCR (BioRad) using ddPCR™ Supermix for Probes (No dUTP) (BioRad cat#1863023) running in BioRad CFX96 Touch Real-Time PCR Detection System. The quality checked libraries were then loaded on a NextSeq 500 running with Nextseq 500/550 high output v2.5 75 cycle kit (Single Read 75 cycles, Cat#: 20024906). The real-time base call (BCL) files were converted to FASTQ files using Illumina bcl2fastq2 (v2) conversion software.

### Analysis of transcriptomics (RNA-seq) data

RNA-seq data processing analysis was carried out using standard procedures and custom R scripts. First, sequenced reads were aligned to the human reference genome GRCh38 and passed through a quality assessment pipeline using the package Rsubread [70]. High read mappability was observed in the dataset and all replicates were included. Next, the mapped reads were counted across all genes using the edgeR R package [71]. Counts-per-million (CPM) values were calculated for all genes to normalize read counts resulting from per-replicate differences of sequencing depths. We focused on transcript abundance values of consensus coding sequence genes (CCDS) database V22 [72] and filtered other classes of genes from our dataset. We also filtered lowly expressed genes and only included genes with above-baseline transcript abundance (CPM > 0.5) in at least two replicates. Next, trimmed mean of M values normalization was performed to remove composition bias between libraries [73]. Two design matrices for comparing the three technical replicates corresponding to distinct siRNAs (H1 and H4, respectively) against the three replicates of the control siRNA (NT1) were generated. Transcript abundance values were subsequently transformed with the voom procedure of the limma package [65]. Differential transcript abundance analysis was conducted by first fitting a linear model to the voom-transformed CPM values. Next, an empirical Bayes shrinkage method was performed on the variances and a statistical test using a pre-defined a fold-change (FC) threshold (abs(log_2_(FC)) > 1.2) was conducted to estimate statistical significance of differential transcript abundance, using the TREAT method [56]. The resulting P-values from the two siRNA experiments (H1 *vs* NT1; H4 *vs* NT1) were merged using the Brown method [74] to prioritize genes differentially regulated in both *HOXA10-AS* depletion experiments and to deprioritize specific off-targets of each of the siRNAs. The merged p-values were corrected for multiple testing using the Benjamini-Hochberg procedure and significant genes were selected (*FDR* < 0.05). To evaluate the agreement of the two siRNAs, we conducted a Pearson correlation test of log10-transformed p-values from the two siRNA experiments. We confirmed that a very small number of significant genes showed opposite fold-changes in the two experiments (3 genes or 0.12%), indicating a strong agreement of the two siRNAs (H1, H4) in depleting *HOXA10-AS* and an overall lack of major off-target effects. Pathway enrichment analysis of differentially expressed genes was conducted using ActivePathways [48] with all genes and corresponding P-values from the two siRNA experiments (H1 *vs* NT1; H4 *vs* NT1) as input and default parameter settings (*FWER* < 0.05). Enrichment maps were generated in Cytoscape using the EnrichmentMap app and standard protocols [47]. Pathway-annotated genes from the ActivePathways analysis were curated for known glioma genes using the COSMIC Cancer Census database [35] and previous GBM sequencing studies [57, 58].

